# An Automatic Glaucoma Grading Method Based on Attention Mechanism and EfficientNet-B3 Network

**DOI:** 10.1101/2023.12.10.571013

**Authors:** Xu Zhang, Fuji Lai, Weisi Chen, Chengyuan Yu

## Abstract

Deep learning has received considerable attention in the computer vision field and has been widely studied, especially in recognizing and diagnosing ophthalmic diseases. Currently, glaucoma recognition algorithms are mostly based on unimodal OCT, the visual field for glaucoma auxiliary diagnosis. Such algorithms have poor robustness and limited help for glaucoma auxiliary diagnosis; therefore, this experiment is proposed to use a 2D fundus image and 3D-OCT scanner two modal data as the experimental dataset and use the EfficientNet-B3 network and ResNet34 network models for feature extraction and fusion to improve automatic glaucoma grading accuracy. Since fundus images usually contain a large number of meaningless black background regions, this may lead to feature redundancy. Therefore, this experiment employs an attention mechanism that focuses the attention of the convolutional neural network on eye subject features to improve the performance of the glaucoma autoclassification model.

## Introduction

Glaucoma, known as the “silent thief of sight”, was recognized as an eye disease in the early 17th century. It begins with damage to the optic nerve due to increased pressure in the eye, called intraocular pressure [1]. The longer the increase in intraocular pressure persists, the more severe the damage to visual function. If left untreated, glaucoma can lead to irreversible visual impairment and even blindness.

According to the latest data, by 2021, approximately 76 million people were suffering from glaucoma worldwide. The number of glaucoma patients in China is approximately 22 million, of which approximately 5.7 million are blind. Therefore, it is imperative to protect your eyesight. Common glaucoma symptoms include sudden loss of vision, severe eye pain, blurred vision, and eye redness. Glaucoma predominantly affects middle-aged and older populations, especially those over the age of 40. Therefore, age is a common risk factor for glaucoma. Early detection and treatment are essential to preventing vision problems caused by glaucoma [2]. Ophthalmologists use different examination methods to examine patients, such as ophthalmoscopy, tonometer, and visual field measurement using a visual field metre for comprehensive glaucoma diagnosis. The ophthalmoscope examination method is used to check the colour and shape of the object, while the tonometer is used to measure internal intraocular pressure. The visual field metre examination is used to analyse the size of the visual field [3]. The complexity of fundus images and the time-consuming and subjective judgement of doctors interfere with traditional manual recognition methods, thus leading to misdiagnosis and omission of glaucoma. Therefore, introducing computer-aided diagnostic (CAD) systems as an important aid to physicians has become particularly urgent. CAD systems play an indispensable role in ensuring an accurate, reliable, and rapid diagnosis of glaucoma [4]. Glaucoma CAD systems can use retinal fundus images as input and classify them as “abnormal” or “normal” by extracting information from multiple feature types. This system can effectively assist doctors in making rapid and accurate glaucoma diagnoses [5]. The Glaucoma CAD system serves a key support role in physician diagnosis, significantly improves physician productivity, reduces physician workload, and greatly reduces the risk of misclassification. The system provides doctors with reliable and objective data, enabling medical teams to process large volumes of fundus images more efficiently and ensuring that patients receive timely and accurate diagnosis and treatment [6]. Artificial intelligence technology can automatically identify glaucoma without much a priori knowledge, so many scholars have attempted to use artificial intelligence technology for patient screening of fundus images. Deep CNNs have proven to be an efficient AI-based tool in identifying clinically significant features from retinal fundus images [7-10].

To obtain a more reliable automatic glaucoma grading algorithm, this project proposes an automatic glaucoma grading method based on the attention mechanism and EfficientNet-B3 network and proposes using two modal data, a 2D fundus image, and a 3D-OCT scanner for model training, testing, and validation to automatically identify glaucoma to achieve higher identification accuracy.

Marcos et al. [11] used a convolutional neural network (CNN) for optic disc segmentation, a step that helps to accurately locate and extract optic disc regions in fundus images. Next, they used advanced image processing techniques to remove nonoptic disc tissues, such as blood vessels, to highlight the features of the optic disc more prominently. The team then focused on extracting textural features of the optic disc region that are important for diagnosing and classifying ophthalmic diseases. Finally, they used these features for classification tasks, which provide reliable support for clinical diagnosis through learning and inference by convolutional neural networks. Raghavendra et al. [12] proposed a novel computer-aided diagnosis (CAD) system for accurately detecting glaucoma using deep learning techniques and designed an 18-layer convolutional neural network. After effective training to extract and then test the features for classification, the system provides a good solution for the early and fast-assisted diagnosis of glaucoma patients. Chai et al. [13] proposed a multibranch neural network (MB-NN) model, which is utilized to adequately extract the deep features from the image and is combined with medical domain knowledge to achieve classification. Balasubramanian et al. [14] attempted feature extraction via a histogram of orientation gradients (HOG) combined with a support vector machine (SVM) for glaucoma classification. However, this method requires tedious preprocessing steps and performs poorly in terms of accuracy. In research on hybrid structures based on structure splicing, Carion et al. [15] proposed DETR, which uses the ResNet backbone network to extract compact image feature representations to generate low-resolution and high-quality feature maps, which effectively reduces the size of the preinput transformer image scale and improves the speed and performance of the model. Ding Pengli et al. [16] proposed CompactNet, a compact neural network based on a compact neural network, to identify and classify retinal images; however, due to the limited experimental samples, the network did not sufficiently extract the relevant features in the training process, so the classification accuracy was not high. Hongjie Gao [17] proposed an algorithm for the vascular segmentation of fundus images with an improved U-shaped network. The algorithm uses the idea of the residual network to change the traditional serial connection method of convolutional layers to the residual mapping phase superposition method and adds batch normalization and a PReLU activation function between the convolutional layers to optimize the network. The algorithm was tested on the DRIVE and CHASE_DB1 fundus databases and compared to the best mainstream algorithms in terms of accuracy, sensitivity, and AUC, which improved by 2.47%, 0.21%, and 0.35%, respectively, on average. Huang Yuankang et al. [18] proposed a method based on Markov random field theory for extracting the optic disc contour of fundus images. Meanwhile, they used the Euclidean distance and correlation coefficient identification method based on the ISNT law for classifying glaucoma fundus images. However, this method requires manual assistance to complete, which is less efficient and less automatic. Panming Li [19] proposed a two-stage automatic SS point localization algorithm based on Gaussian heatmap regression and deep reinforcement learning. Optimal classification performance was achieved in a classification model based on SE-ResNet18.

Generally, at present, domestic and foreign research teams mainly use traditional neural networks for glaucoma recognition research, which has the following defects. First, its dataset only adopts the most common 2D fundus image with a single modality. Second, it fails to account for the large and meaningless black background in the 2D fundus image, which makes its performance poor. Finally, the above methods have scope for improvement in terms of accuracy, kappa value, recall, and F1 value. Rate and F1 value have room for improvement. However, this experimental method adopts two modal data, a 2D fundus colour photo and a 3D OCT scanner, as the experimental dataset, which achieves multimodality and more accurate image feature extraction. Second, this experiment fully analyses the characteristics of the fundus images and uses the attention mechanism to discard the meaningless black background so that the convolutional neural network focuses more on the main features of the eye, thus improving the recognition and grading performance. Moreover, this experimental goal is to achieve higher accuracy, kappa value, recall, and F1 value, so the use of the EfficientNet-B3 network, which is relatively new, can more accurately achieve automatic glaucoma grading.

## Data and Methodology

### 2.1 Dataset

The dataset used in this experiment was provided by Zhongshan Ophthalmology Centre, Sun Yat-sen University, Guangzhou, China, which contains 200 data pairs of two clinical modality images: 100 pairs in the training set and 100 pairs in the test set. The two modalities are 2D fundus image colour photographs and 3D optical coherence tomography (OCT), commonly performed in clinical fundus examinations. For deep learning algorithms, 100 training data pairs are small samples, so the recognition model proposed in this experiment is suitable for training on small sample datasets. Figure 1 shows some 2D fundus images, which will be used for EfficientNet-B3 network training. Figure 2 shows some 3D-OCT images, which will be used for ResNet34 network training after convolution.

**Figure 1:**
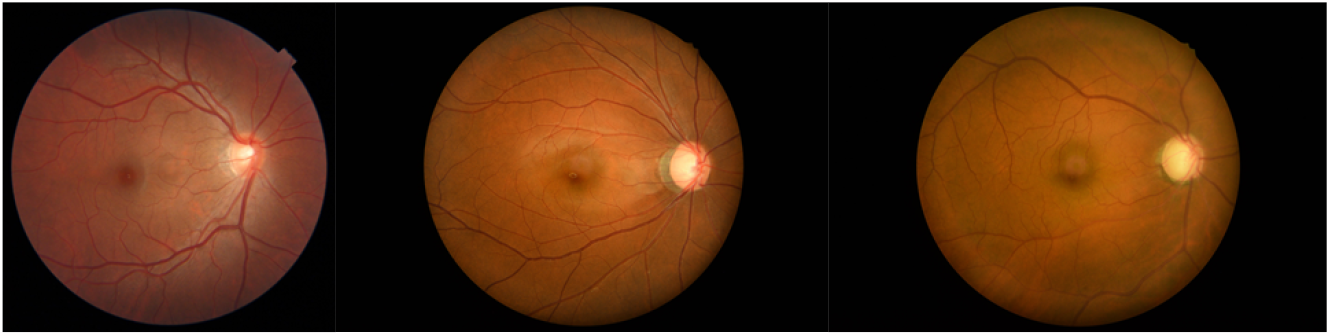
2D fundus images

**Figure 2:**
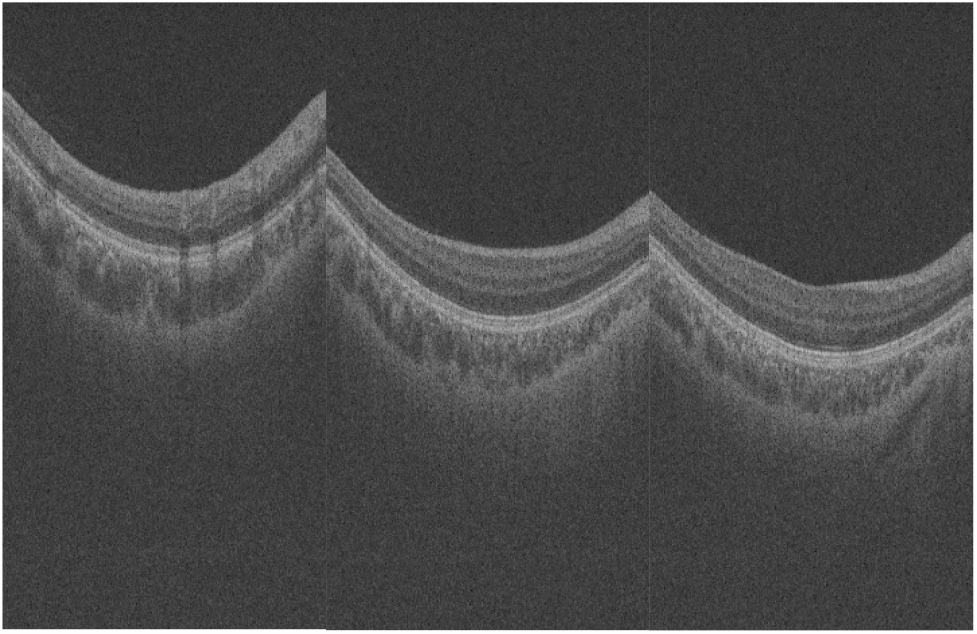
3D-OCT scanner images

### 2.2 Data Preprocessing

In this experiment, data enhancement methods such as flipping (horizontal + vertical), adding noise, random rotation, random flip, random change in brightness, random change in contrast, random change in saturation, cropping, scaling/stretching, and blurring are used. By applying these approaches individually or in combination, the dataset can be processed to increase the quantity of data, capture more image features, and enable the model to see more data variations, improving the model’s generalizability. The use of these data enhancement modalities can effectively address the lack of data volume and mitigate the overfitting problem of the model, as well as improve the model’s ability to adapt to new data, which can help enhance the model’s ability to perform in the glaucoma autoclassification task.

### 2.3 EfficientNet-B3 Network Training Model

EfficientNet-B3 is a network model with unique features whose design benefits from the experience of other good neural networks. The network model contains a residual structure that not only deepens the depth of the network but also makes feature extraction more accurate and efficient. In addition, it allows the flexibility of adjusting the number of feature layers in each layer to achieve more layers of feature extraction, thus enhancing the width of the network. In addition, EfficientNet-B3 can learn and express information from richer data by enlarging the input image solution resolution, which can help enhance model precision. Overall, EfficientNet-B3 is an efficient and flexible network model that draws on the design of many excellent neural networks, making it perform well in a variety of tasks. Figure 3 is a schematic structural diagram of the EfficientNet-B3 network.

**Figure 3:**
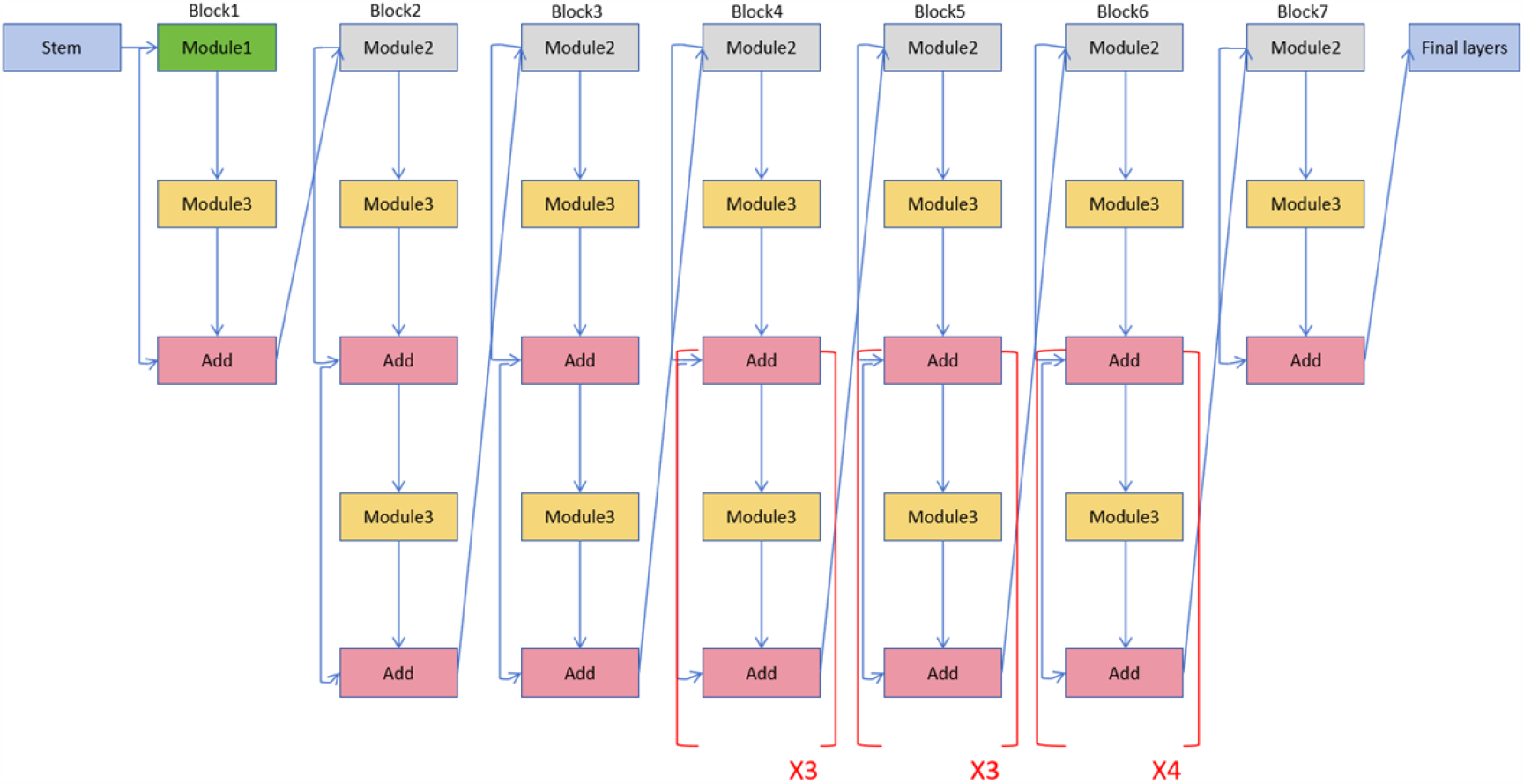
EfficientNet-B3 network structure

### 2.4 ResNet34 Network Model

In deep learning, deep neural networks are a very effective model. However, as the network layers increase, some problems arise, such as gradient vanishing and gradient explosion. These problems make training deep networks very difficult. Therefore, to overcome these problems, there are some solutions, one of which is ResNet, which is a kind of residual neural network. The fundamental concept is to construct a deep network by adding “residual blocks”. The core idea is to introduce cross-layer connections in the network so that the information can be passed directly from the front layer to the back layer. Such cross-layer connections can relieve the gradient vanishing and gradient exploding problems effectively, making training deep networks easier. With ResNet, networks can become deeper without causing performance degradation or training difficulties. This makes ResNet an essential breakthrough in the deep learning field, helping to solve more complex tasks and handle larger data. As a result, ResNet has been widely used in deep learning research and practical applications.

ResNet34 is a relatively concise ResNet structure containing 34 convolutional layers and 18 residual blocks. It is inspired by solving the vanishing and exploding gradient problems in deep neural networks. First, the input layer, ResNet34, is an ordinary convolutional layer including 64 convolutional kernels, each of which is 7×7 in size, with a stride of 2 and padding of 3. The main purpose of this layer is to cut the input image in half in terms of size and to extend the lower-level features out of the image. Next is the residual block. ResNet34 has a total of 18 residual blocks. Every residual block is composed of two convolutional layers of size 3 × 3 and a cross-layer connection. For the first convolutional layer, the step size is 1, and the padding is 1. The second convolutional layer also has a step size of 1 and a padding of 1. The cross-layer connection enables the outer output of the former layer to be directly appended to the input of the latter layer, which preserves the message of the earlier layer and passes it on to the latter layer. This design helps in information transfer and mitigates the vanishing and exploding gradient problems. Thus, ResNet34 effectively addresses some of the difficulties in training deep neural networks by introducing residual blocks and cross-layer connections. This design allows the network to be deeper, easier to train and to achieve excellent performance with relatively few parameters. Furthermore, there is a global average pooling layer, which is added following the latest residual block of ResNet34. The role of this layer is to average pool the output of the last residual block to obtain a global feature. Global average pooling is an operation that compresses the entire feature map into a single value. Through this operation, the network can obtain comprehensive information about the whole image and thus better understand the overall semantics. Finally, there is a full-connectivity layer, which is inserted following the global average pooling layer. The role of this layer is to map global features to category scores. The full-connectivity layer is typically used for performing classification tasks, where associations are established between the extracted features and the categories to obtain scores or probabilities for different categories. Ultimately, the model makes classification decisions based on these scores or probabilities. With the global average pooling layer and the fully connected layer, ResNet34 can map image features to the final category scores to perform tasks such as image classification. Introducing global average pooling and fully connected layers gives ResNet34 a powerful classification capability and allows it to perform well in a variety of image recognition problems. Figure 4 is a schematic diagram of the Residual block structure.

**Figure 4:**
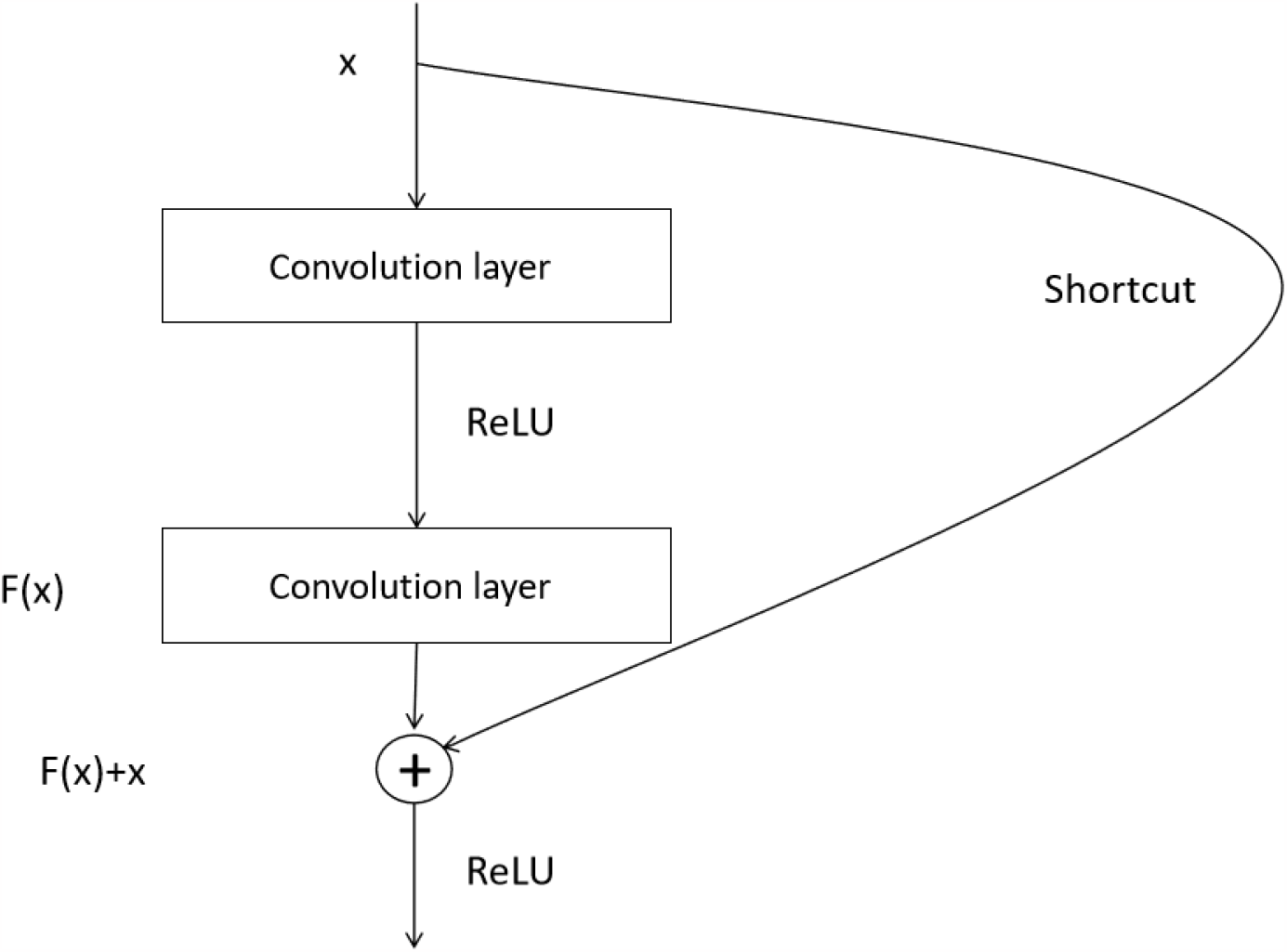
Residual block

### 2.5 Attention Mechanism

Attentional mechanisms have achieved significant success in computer vision tasks but have been less frequently applied in glaucoma recognition and automatic grading tasks. This may be because the job involves complex medical images, small datasets, and difficult labelling, while the attention mechanism requires larger datasets and expert support. However, applying attention mechanisms in glaucoma recognition and automatic grading is still worth exploring as profound learning techniques advance and datasets increase.

The attentional mechanism works by focusing on the most salient parts of the characteristics that are extracted by the deep neural network, thereby eliminating redundant information from the visual task. This mechanism is usually implemented by embedding an attention map into the neural network. The attention mechanism allows the neural network to automatically learn and select the most critical regions and features in the image while ignoring the unimportant parts. In this way, the network can focus on meaningful information more effectively, improving task performance and efficiency. The fundus image in this experimental dataset contains many pointless black background areas, which can result in many redundant features, so this experiment adds an attention mechanism to the feature extraction process so that the convolutional neural network focuses more on the main features of the eyes, thus improving the performance of the glaucoma autoclassification model.

### 2.6 Loss Function

Since automatic glaucoma classification is a three-classification task, the softmax function plus the cross-entropy loss function is adopted as the loss function in this paper. Using the softmax function can restrict the output between 0 and 1, and the sum of the probabilities of each sample falling into each category is just 1. The cross-entropy loss value is calculated by the formula: 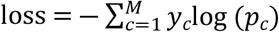 where M is the number of categories, *y*_*c*_ is equal to 0 or 1, 1 if the predicted category is the same as the sample labelling, 0 otherwise, and *p*_*c*_ is the probability that the sample belongs to category c.

Finally, the loss results of all the samples in the training set are summed to obtain the final total loss.

## Results

### 3.1 Experimental Environment

All the algorithms in this article are performed in the following hardware environment. GPU: TeslaV100, Memory: 32GB, CPU: 4 cores, implemented using the Python language based on the paddle deep learning framework.

### 3.2 Results and Analyses

To validate the usefulness of the data extension method, in this paper, we trained the EfficientNet-B3 model on the primary dataset and the extended dataset for 50 epochs. After training was completed, we evaluated the model using an independent test set and recorded the accuracy of the model. By comparing the performance of the models trained on the original and expanded datasets, we can conclude that the data expansion method has a positive effect on model performance.

Table 1 demonstrates the effect of data expansion on model performance. The accuracy of the EfficientNet-B3 model on the original dataset is 96.61%, whereas, through data expansion, the model accuracy is increased to 97.58%, which is an improvement of 0.97%. This result shows that model accuracy and performance are significantly enhanced by data enlargement.

**Table 1:** Comparison of the test results before and after expanding the data.

| Dataset | Model | Epoch | Accuracy |
| --- | --- | --- | --- |
| Primary dataset | EfficientNet-B3 | 50 | 96.61% |
| Augment dataset | EfficientNet-B3 | 50 | 97.58% |

To continue to enhance the model’s efficiency, this paper adopts an effective method for removing unnecessary redundant features in the convolutional neural network by applying an attention mechanism to the EfficientNet-B3 network structure, which can significantly improve the performance of glaucoma autograding. GoogLeNet [20], for the first time, proposed the use of convolutional kernels of multiple sizes for simultaneous feature extraction, and this method is known as the inception module. By introducing the inception module, GoogLeNet can extract a wide range of features at different scales and levels, thus increasing the network width and allowing it to better capture information at different scales. However, ResNet has successfully strengthened the delivery of gradient information across the network through feature reuse. This result is due to the introduction of residual blocks and cross-layer connectivity. This design allows information to be passed in jumps, mitigating the problems of vanishing and exploding gradients while also allowing the network to become deeper and easier to train. These two network architectures have excellent performance in many computer vision tasks, so GoogLeNet and ResNet are constructed as baseline models for comparison in this experimental work. In this experiment, GoogLeNet, ResNet, and the presented model are tested on the extended dataset with 200 training epochs. The tested model is confirmed on the test set, and the accuracy, kappa value, recall, and F1 value of the model are calculated. The results of the tests are displayed in Table 2.

**Table 2:** Comparing the results of tests with the baseline model.

| Model | Accuracy | kappa | Recall | F1-score |
| --- | --- | --- | --- | --- |
| GoogLeNet | 94.79% | 0.9714 | 94.32% | 95.91% |
| ResNet | 96.84% | 0.9802 | 96.69% | 96.11% |
| EfficientNet-B3+ResNet34 | **97.83%** | **0.9911** | **97.24%** | **97.03%** |

Table 2 demonstrates the results of the trials for the baseline models, in which the accuracy, kappa value, recall, and F1 values are 94.79%, 0.9714, 94.32%, and 95.91% for GoogLeNet and 96.84%, 0.9802, 96.69%, and 96.11% for ResNet, whereas the values of accuracy, kappa value, recall and F1 value for EfficientNet-B3+ResNet34 are 97.83%, 0.9911, 97.24% and 97.03%, respectively.

compared with the baseline model, the accuracy, kappa value, recall, and F1 value of the EfficientNet-B3+ResNet34 proposed in this paper, which is built on the attention mechanism, are at the optimal level in all four metrics, so it can be demonstrated that by using both modal data of 2D and 3D OCT scanners as the experimental dataset and using the EfficientNet-B3+ResNet34 network for glaucoma recognition and autograding, not only is the accuracy improved but its performance is also enhanced.

### 3.3 Comparison with the Literature

To further highlight the value and contribution of the work in this article, the proposed model is analysed in comparison with the existing achievements in the literature. The results of the comparison are detailed in Table 3 below.

**Table 3:** Comparative results with the literature.

| Method | Model | Accuracy |
| --- | --- | --- |
| Balasubramanian et al<sup>[29]</sup> | HOG | 96.08% |
| Ding Pong Li et al. | CompactNet | 97.60% |
| Gao Hongjie et al. | Improved U-shaped network | 97.69% |
| Proposed method | EfficientNet-B3+ResNet34 | **97.83%** |

Table 3 gives the results of the comparison between the work in this paper and the work in the literature, in which Balasubramanian et al. achieved an accuracy of 96.08% in glaucoma classification using histogram of orientation gradients (HOG) for feature extraction and combining it with support vector machines (SVMs), while Pongli Ding et al. used a compact neural network based on CompactNet, with an accuracy of 97.60%. Hongjie Gao et al. used an improved U-shaped network with an accuracy of 97.69%. In this paper, after the data enlargement process, the EfficientNet-B3+ResNet34 for the attention-based regime was intensively trained, and a model with excellent performance was obtained. On an independent testing set, the model demonstrated an accuracy as high as 97.83%, which fully proved the value of the model in the field of glaucoma automatic identification.

By comparing the current model with the most accurate glaucoma recognition rate, the accuracy of this experimental model for recognition and automatic grading is better than the current latest, most effective method, therefore demonstrating the validity of the work in this article.

## Conclusion

In this paper, for the problem of lacking data samples, the data enhancement method is used to expand each sample in the dataset, experiments are carried out, and the experimental results validate the validity of the extension of the data for automatically grading glaucoma. For the large pointless black background areas in each fundus image, this paper proposes the method of introducing the attention module to allow the convolutional neural network to focus more on the main structure of the eye in the image and extract the key features to enhance the efficiency of the glaucoma autoclassification model. In contrast, the approach in this article is evaluated against the baseline model and relevant research, and the experimental results indicate that the method suggested in this article can significantly increase the accuracy of automatic glaucoma grading and reach the optimum in all aspects. In the future, based on ensuring the model properties, we will further study the lightweight model structure to achieve actual model deployment and use it in clinically assisted diagnosis and treatment.

## Acknowledgement

The authors would like to thank the reviewers and editors for their important and helpful comments, which greatly improved the quality of this paper.

